# The A domain of clonal complex 1-type fibronectin binding protein B promotes adherence and biofilm formation in *Staphylococcus aureus*

**DOI:** 10.1101/2023.04.25.538241

**Authors:** Sara R. Henderson, Joan A. Geoghegan

## Abstract

Adhesive interactions between *Staphylococcus aureus* and the host rely on cell wall-anchored proteins such as fibronectin binding protein B (FnBPB). Recently we showed that the FnBPB protein expressed by clonal complex (CC) 1 isolates of *S. aureus* mediates bacterial adhesion to corneodesmosin. The proposed ligand binding region of CC1-type FnBPB shares just 60% amino acid identity with the archetypal FnBPB protein from CC8. Here we investigated ligand binding and biofilm formation by CC1-type FnBPB. We found that the A domain of FnBPB binds to fibrinogen and corneodesmosin and identified residues within the hydrophobic ligand trench in the A domain that are essential for the binding of CC1-type FnBPB to ligands and during biofilm formation. We further investigated the interplay between different ligands and the influence of ligand binding on biofilm formation. Overall, our study provides new insights into the requirements for CC1-type FnBPB-mediated adhesion to host proteins and FnBPB-mediated biofilm formation in *S. aureus*.

## Introduction

*Staphylococcus aureus* is a pathogen of humans capable of causing a variety of infection types. It is also a commensal that colonises multiple body sites including the nares of healthy individuals asymptomatically and skin of atopic dermatitis (AD) patients. Colonisation and the development of infection relies on adhesive interactions between *S. aureus* and host proteins and cells. Microbial Surface Components Recognising Adhesive Matrix Molecules (MSCRAMM) proteins such as fibronectin binding protein B (FnBPB) are anchored to the cell wall of *S. aureus* and displayed at the cell surface. They play a key role in mediating adhesion. MSCRAMMs share a conserved domain architecture (Fig. 1A). The best characterised MSCRAMMs, ClfA and SdrG, bind their ligands through a mechanism known as “dock, lock and latch” utilising a ligand binding trench that is formed between the independently folded N2 and N3 domains (1, 2).

**Fig. 1.**
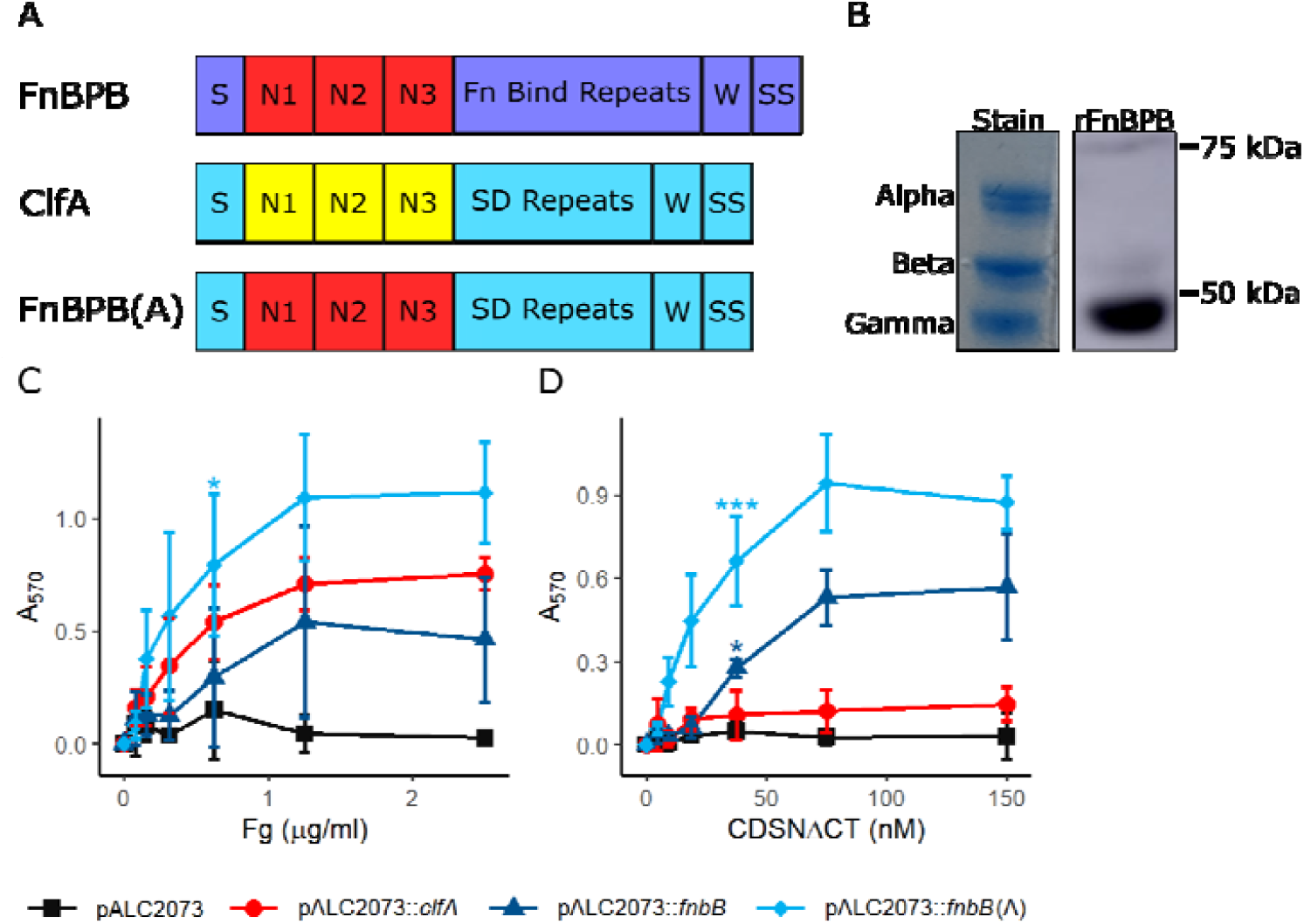
The role of the CC1-type FnBPB A domain in ligand adhesion. **A)** Schematic of the domain organisation of FnBPB, ClfA and FnBPB(A) proteins. The signal sequence (S) is followed by the N1N2N3 domains (A region) (red CC1-type FnBPB, yellow ClfA), either a series of fibronectin (Fn) binding repeats (FnBPB) or serine-aspartate (SD) repeats (ClfA, FnBPB(A)) which form the stalk that connects the N2N3 domains to the surface of the bacterium. The wall spanning region (W) is linked to the sorting signal (SS) which allows for processing of the proteins by sortase A to allow the C-terminus of the protein to be anchored to the peptidoglycan. B) Fibrinogen was separated on 10% Bis-tris Gels in MOPS running buffer under reducing conditions. The stained portion of the gel (stain) was used to distinguish the positions of the alpha, beta and gamma chains of fibrinogen. The rFnBPB portion of the gel was transferred to PVDF membrane, blocked with BSA (5% w/v) overnight (4°C), probed with recombinant rFnBPB for 4 h at room temperature. Bound rFnBPB was detected with HRP-conjugated Anti-His antibody (Roche, 1:1000), and incubated with LumiGlo reagent before chemiluminescent imaging. Adhesion of SH1000 Δ*clfAclfBfnbAfnbB* to C) fibrinogen or D) CDSNΔCT. Black *Square* pALC2073, red *Circle* pALC2073::*clfA*, Blue *Triangle* pALC2073::*fnbB*, Pale blue *diamond * pALC2073::*fnbB*(A). Data is presented as the mean of a minimum of 3 biological replicates, error bars represent ±1 standard deviation of the mean. Statistical significance determined by 1-way ANOVA at 0.625 μg/ml fibrinogen (B) or 37.5 nM CDSNΔCT (C) with post-hoc Dunnett’s test of multiplicity against the pALC2073 strain. * P<0.05, ** P<0.01, *** P<0.001

FnBPB plays a key role in promoting the adhesion of *S. aureus* to corneocytes, cells present in the uppermost layer of the stratum corneum (3). This is believed to be mediated by its interaction with corneodesmosin (CDSN), a protein that localises to the tips of villus like projections that protrude from the surface of corneocytes as a result of loss of natural moisturising factor from the corneocyte in subjects with moderate to severe AD (4, 5). Studies to date of the interactions between *S. aureus* and corneocytes in AD have mainly focused on an *S. aureus* strain from clonal complex (CC)1 (3) due to this being the most prevalent CC isolated from people with AD in certain geographical locations (6–8). The FnBPB protein expressed by CC1 *S. aureus* (CC1-type FnBPB) shares just 60% amino acid identity in the N2 and N3 domains to the archetypal FnBPB protein produced by CC8 strains. The N2N3 domains of region A of the CC8-type FnBPB bind to fibrinogen, plasminogen, elastin and histones (9–11). FnBPB from different CCs bind to histones with different affinities, suggesting that sequence variants may have differences in their ligand binding specificity (9). At present there are no structures of the FnBPB N2N3 domains from any clonal complex.

Previously, mutagenesis revealed that a N312A and F314A substitutions in the N2 domain of CC8-type FnBPB resulted in a significant loss of fibrinogen (Fg) binding but not plasminogen binding (12, 13). Likewise, in other MSCRAMMs it has been shown that residues at the equivalent position to N312 result in a significant loss of Fg binding while mutations at the position of F314 result in a complete loss of Fg binding comparable to the double mutation (2, 10, 13). The stalk region of FnBPB and the related protein FnBPA comprises a series of fibronectin binding repeats (FnBRs) (14). As well as promoting adhesion to fibronectin, amino-acid sequence variation in the FnBRs of FnBPA has been demonstrated to play an important role in modulating the binding of fibrinogen to FnBPA (15).

In additional to their roles in adhesion, FnBPA and FnBPB promote the accumulation phases of biofilm formation in *S. aureus*. It has been shown in clonal complexes 8, 22 and 30 that biofilm formation by methicillin resistant *S. aureus* grown in medium supplemented with glucose is dependent on the presence of FnBPA or FnBPB (16–18). Biofilm accumulation was shown to be polysaccharide independent and mediated by protein-protein interactions involving the A region, but independent of the ligand binding trench of FnBPA (16, 19). Clinical isolates from skin and soft tissue infections formed enhanced biofilms in the presence of glucose and fibrin suggesting that fibrinogen or it’s break down product fibrin may be incorporated into biofilms (20).

Here we sought to characterise region A of CC1-type FnBPB in terms of its ligand binding activity and ability to mediate biofilm formation in *S. aureus*. To do this we expressed CC1-type FnBPB and mutants deficient in adhesion and biofilm formation in *S. aureus*. Our findings shed new light on the mechanism by which adhesion occurs, and how this protein may support the formation of a biofilm, potentially contribution to skin colonisation and infection.

## Results

### The A domain of CC1-type FnBPB mediates bacterial adhesion to corneodesmosin and fibrinogen

To understand more about the interaction of CC1-type FnBPB with ligands we studied a mutant of *S. aureus* deficient in ClfA, ClfB, FnBPA and FnBPB (SH1000 *clfAclfBfnbAfnbB*) carrying full-length CC1-type FnBPB on a plasmid (pALC2073::*fnbB*). SH1000 *clfAclfBfnbAfnbB* (pALC2073::*fnbB*) adhered to Fg in a dose-dependent manner while SH1000 *clfAclfBfnbAfnbB* carrying an empty vector (pALC2073) did not (Fig. 1C). To explore if the N-terminal A region of FnBPB (domains N1,N2,N3) is sufficient to support bacterial adhesion to Fg, a hybrid protein where the A region of FnBPB was fused to the stalk region of ClfA containing serine aspartate (SD) repeat domains rather than the fibronectin binding repeats (Fig. 1A), was expressed on the surface of *S. aureus* from plasmid pALC2073::*fnbB*(A). The SD repeat domains were previously shown to lack ligand binding activity (21). The hybrid protein adhered to Fg demonstrating that the N-terminal A region of FnBPB is sufficient to support adhesion (Fig. 1C). Bacteria expressing ClfA also bound to Fg as expected given that the A region of ClfA binds to fibrinogen (21). Previously the N2N3 domains of FnBPA, FnBPB (CC8-type) and ClfA have all been shown to bind to the extreme C-terminus of the γ-chain of Fg (13, 22, 23) while ClfB, which like CC1-type FnBPB bind to CDSN and loricrin (3, 24, 25), binds to the α-chain of fibrinogen (26). To determine which chain the A domain of CC1-type FnBPB binds to, we separated Fg by denaturing SDS-PAGE before performing far-western blotting analysis with CC1-type recombinant FnBPB N2N3. Our results clearly show that CC1-type FnBPB binds only to the γ-chain of fibrinogen, and we were unable to detect an interaction with the other chains (Fig. 1B). Therefore CC1-type FnBPB appears to bind specifically to the Fg γ-chain, like ClfA and FnBPA.

The N-terminal glycine-serine rich region of CDSN comprises the FnBPB binding site (3). Here, bacterial adherence to a GST-tagged truncate of CDSN containing the entire FnBPB binding site but lacking the C-terminal glycine-serine rich loop region (CDSNΔCT), was examined. Bacteria expressing CC1-type FnBPB or the hybrid protein [SH1000 *clfAclfBfnbAfnbB* pALC2073::*fnbB*(A)] adhered to CDSNΔCT (Fig. 1D) while bacteria carrying empty vector or expressing full-length ClfA did not, demonstrating that both the stalk region, consisting of SD repeats, and the Fg-binding A region of ClfA do not support adherence to CDSNΔCT. Adhesion to GST alone was negligible for all strains indicating that this assay is specifically detecting interactions with CDSNΔCT (Fig. S1A). Bacteria expressing hybrid protein appeared to adhere to CDSNΔCT to at a higher level than bacteria expressing full-length FnBPB suggesting either that the A domain is more exposed in the hybrid protein, or that this protein is expressed at higher levels than FnBPB. In summary these results demonstrate that the CDSN binding site is contained within the A region of CC1-type FnBPB and that the FnBPB A domain binds to the γ-chain of Fg.

### FnBPB binds to Fg and CDSN through the hydrophobic trench between domains N2 and N3

Next, we set out to identify residues in CC1-type FnBPB that are required for binding to CDSN and Fg. The N2N3 domains of CC1-type FnBPB protein shares 60% amino-acid identity with the previously characterised CC8-type FnBPB and 50% identity with FnBPA (Fig. S2). The crystal structure of CC8-type FnBPA has been solved in complex with a peptide comprising the C-terminal residues of the γ-chain of fibrinogen allowing the contact residues to be identified (27). Amino-acid substitution of residues N306 and F308 in FnBPA abolished binding of recombinant N2N3 domain protein to fibrinogen (10). In the absence of a crystal structure of FnBPB we made sequence alignments and structural predictions and identified N277 and F279 as adopting the equivalent positions in CC1-type FnBPB (Fig. S2) (28–30). We investigated if substituting N277 and F279 with alanine, either singly or in combination, reduced the ability of CC1-type FnBPB to promote adherence to Fg and CDSNΔCT. Bacteria expressing the CC1-type FnBPB A region with an N277A substitution had reduced adherence to Fg (Fig. 2A) and CDSNΔCT (Fig. 2B) compared to the control carrying wild-type sequence. The protein harbouring an N279A or N277A+F279A substitutions did not support bacterial adherence to Fg or CDSNΔCT (Fig. 2). We confirmed that the variant proteins harbouring amino-acid substitutions were expressed at similar levels to their parent protein (Fig. S3) indicating that the differences in ligand binding were likely due to disruption of the ligand binding trench (Fig. S3). Additionally, far-western blotting with rFnBPB N277A/F279A showed weak interaction with the γ-chain of Fg (Fig. S4). Together these results indicate that both N277 and F279 are involved in the interaction of CC1-type FnBPB with CDSN and Fg, with F279 potentially being more important for the binding.

**Fig. 2.**
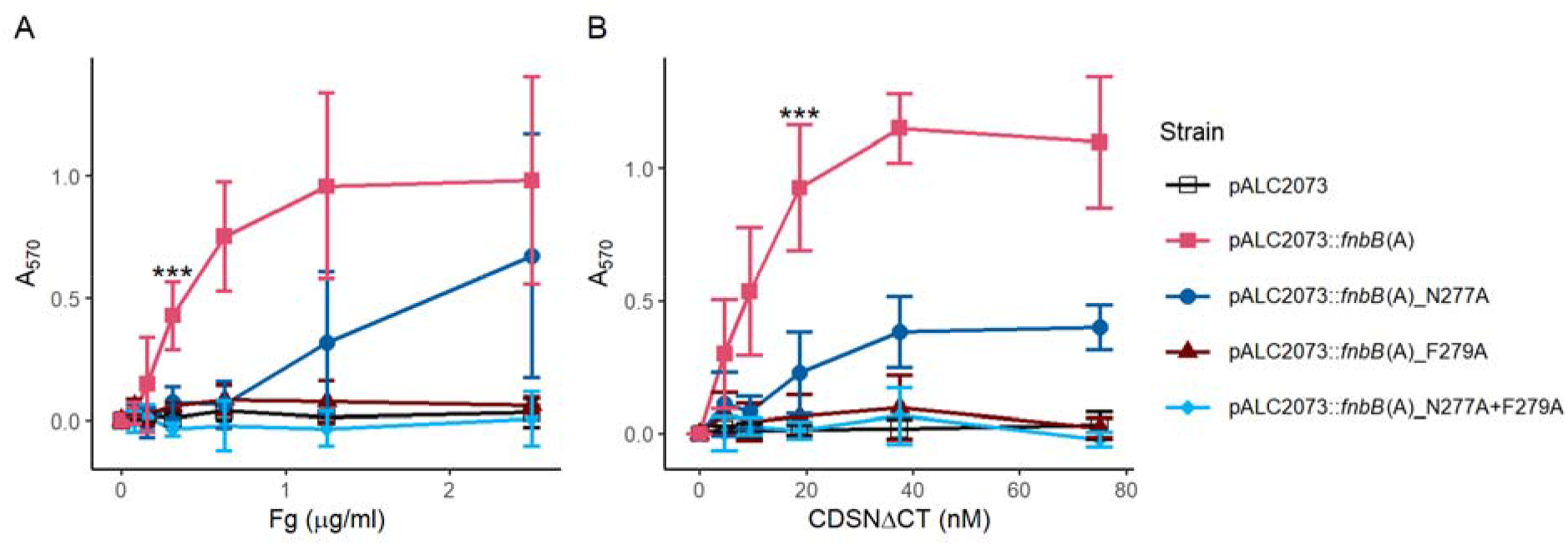
FnBPB mediates ligand binding through the hydrophobic trench between the N2 and N3 domains. Adhesion of SH1000 *clfAclfBfnbAfnbB* carrying the pALC2073::*fnbB*(A) plasmid with mutations in the hydrophobic trench between N2 and N3 to A) fibrinogen or B) corneodesmosin (CDSNΔCT). Data is presented as the mean of a minimum of 3 biological replicates, error bars represent 1 standard deviation from the mean. Statistical analysis via 1-way ANOVA at A) 0.3125 μg/ml Fg or B) 18.75 nM CDSNΔCT with a post-hoc Dunnett’s test of multiplicity against the wild type pALC2073::*fnbB*(A) strain. *** P<0.001.

### Fibrinogen and corneodesmosin partially inhibit FnBPB-mediated adherence to immobilised ligands

Next, we were interested to see if corneodesmosin and fibrinogen compete for binding to the FnBPB A domain, or if the protein could bind to both ligands at the same time. To determine this, we carried out a competition assay whereby bacteria expressing pALC2073::*fnbB*(A) were preincubated with free ligands before being added to wells coated in Fg or CDSNΔCT. The results showed that the proportion of cells adhering to immobilised Fg and CDSNΔCT was reduced by approximately 35 - 60% by preincubation with either Fg or CDSNΔCT. Free ligand concentrations 10 times higher than the immobilised ligand concentration were unable to completely abolish adhesion to either immobilised ligand. Bacterial adhesion to Fg and CDSNΔCT was unaffected by incubation with GST. As a control we showed that the bacteria did not bind to immobilised GST under any of the conditions (Fig. 3). In conclusions these data indicate that adherence to immobilised Fg and CDSNΔCT can be partially inhibited by preincubation with either Fg or CDSNΔCT and suggests that these ligands may interact with overlapping binding sites in CC1-type FnBPB.

**Fig. 3.**
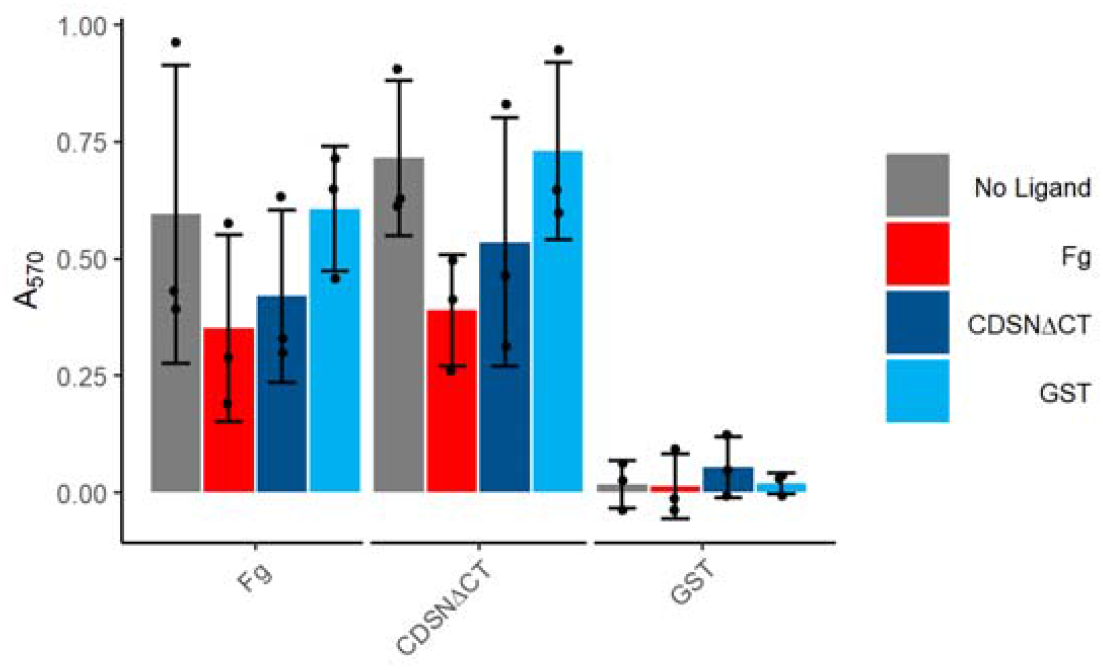
Fibrinogen and corneodesmosin bind competitively to the FnBPB A domain. SH1000 *clfAclfBfnbAfnbB* expressing pALC2073::*fnbB*(A) grown to an OD_600_ =0.35 were preincubated with Fg (8 μg/ml) or GST-CDSNΔCT (750 nM) or GST (750 nM) or PBS (no ligand) for 30 min before addition to plates containing immobilised Fg (0.8 μg/ml), CDSNΔCT (75 nM), GST (75 nM). Data is presented as the mean of a minimum of 3 biological replicates, error bars represent 1 standard deviation of the mean. No significance was found using a 1-way ANOVA with Tukey’s honest significance test.

### Region A of CC1-type FnBPB mediates biofilm formation

Previously it was shown that CC8-type FnBPB and FnBPA mediate biofilm accumulation in *S. aureus*. The A region of FnBPA is required to support biofilm formation, but the fibronectin binding repeats are not (18). FnBPA- and FnBPB-mediated biofilm formation require mildly acidic growth conditions which are achieved by growing the bacteria in medium supplemented with glucose (16). Hence, we explored if CC1-type FnBPB mediates biofilm formation in TSB supplemented with glucose (TSBG). The hybrid protein (FnBPB region A fused to the SD repeat region of ClfA) supported biofilm formation when the growth medium was supplemented with anhydrotetracycline to induce expression of the gene under the control of the promoter on pALC2073 (Fig. 4A). No biofilm formation occurred in the absence of inducer (Fig. 4A). In contrast, little to no biofilm was formed by strains expressing *clfA* (Fig. 4A). This suggests that region A of FnBPB (CC1-type) is capable of supporting biofilm formation. Surprisingly, bacteria expressing full-length *fnbB* did not form biofilm (Fig. 4A).

**Fig. 4.**
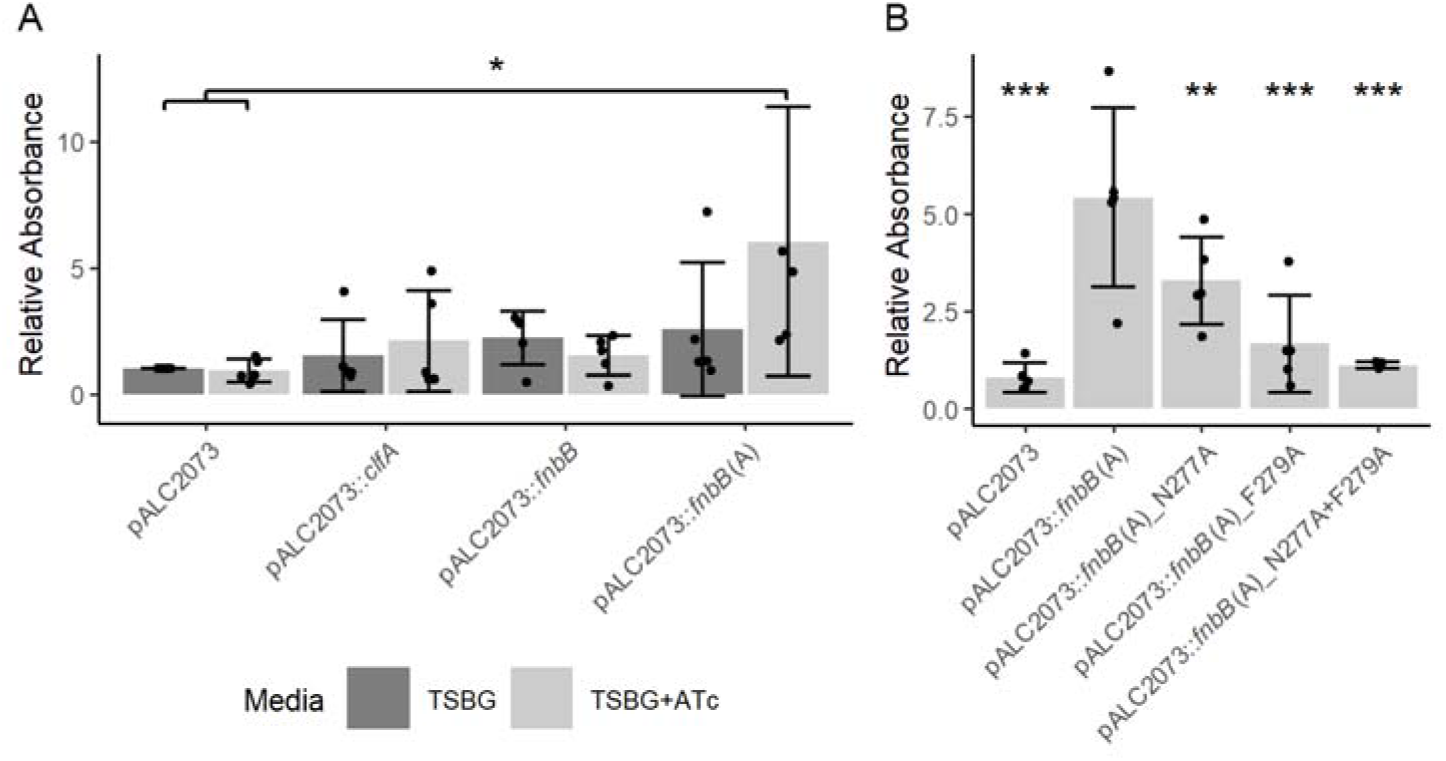
The role of the FnBPB A domain in biofilm formation. Biofilm formation of SH1000 *clfAclfBfnbAfnbB* transformed with A) plasmids expressing the *clfA, fnbB* or *fnbB*(A) genes or B) plasmids expressing the *fnbB*(A) mutant variants. Bacteria were grown in tryptic soy broth supplemented with glucose (TSBG) and the absorbance was expressed relative to the empty vector in the absence of inducer. ATc= biofilm grown in the presence of anhydrotetracycline (150 ng/ml). Biofilms were washed and subsequently quantified by staining with crystal violet (1% w/v) solubilised in 70% ethanol. To determine the relative absorbance values the A_570_ values for each biological replicate were divided by those of pALC2073 in TSBG. Data is presented as the mean of 5 biological replicates, error bars represent 1 Standard deviation from the mean. Error bars represent 1 standard deviation of the mean. Statistical analysis via 1-way Anova with A) Tukey’s honest significance test or B) Dunnett’s Multiple comparison test relative to the pALC2073::*fnbB*(A). *p<0.05, **p<0.01, ***p<0.001

Previously, it was found that an FnBPA residue involved in binding to Fg (N304A) (18) was not required for promoting biofilm formation. We therefore examined if the amino-acid substitutions in the CC1-type FnBPB A domain that reduce adhesion to Fg and CDSN affect biofilm formation. Bacteria expressing the hybrid protein harbouring an N277A substitution in region A formed significantly less biofilm than those expressing the hybrid protein with the native FnBPB region A sequence. The proteins harbouring amino-acid substitutions F279A or N277A+F279A showed a greater reduction in biofilm formation (Fig. 4B).

### Biofilm formation is not inhibited by the addition of FnBPB ligands

As adhesion to ligands and biofilm formation were both inhibited by the same amino-acid substitutions in the ligand binding trench, we were interested to see if ligand binding inhibited biofilm formation. Our results showed that biofilm formation by SH1000 *clfAclfBfnbAfnbB* (pALC2073) was unaffected by the inclusion of ligands in the growth medium (Fig. S5). We determined that a non-significant but apparently concentration dependent increase in biofilm formation occurred on addition of CDSNΔCT or Fg to the media (Fig. 5). Ligands did not alter biofilm formation for either the full-length FnBPB or the FnBPB A region mutant deficient in ligand binding (Fig. S5). In the presence of Fg a biofilm was produced in a ClfA-expressing strain (Fig. S5). This demonstrates that the mechanism of biofilm formation is complementary to ligand binding and that the ligands may act as bridges between bacteria within the biofilm.

**Fig. 5.**
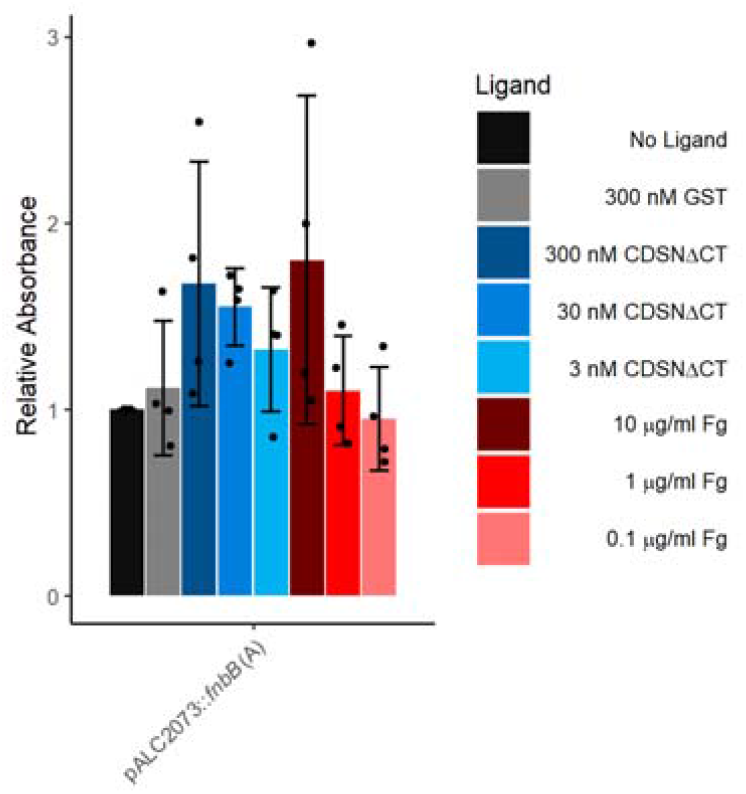
The impact of FnBPB ligands on biofilm formation. Biofilms were grown in TSBG with anhydrotetracycline (150 ng/ml) with ligands added at the specified concentrations. Biofilms were washed and subsequently quantified by staining with crystal violet solubilised in 70% ethanol. To determine the relative absorbance values the A570 values for each biological replicate were divided by those obtained in the absence of a ligand. Data is presented as the mean of 4 biological replicates. Error bars represent 1 standard deviation from the mean. No significance was found by performing a 1-way ANOVA test with a Dunnett’s Multiple Comparison test comparing each strain to the biofilm formed in the absence of any ligand.

## Discussion

We recently discovered that FnBPB promotes adherence of *S. aureus* to the surface of corneocytes during the colonisation of atopic dermatitis skin (3). The N-terminal region of CDSN was shown to be the ligand. However, the binding mechanism was not investigated. Here we demonstrate that the FnBPB A domain mediates binding to CDSN and also to fibrinogen and that residues N277 and F279, predicted to reside in the hydrophobic trench between the N2 and N3 domains are important for interaction with ligands. The results indicate that phenylalanine at position 279 is more critical in ligand binding than asparagine at position 277. This agrees with previous studies showing that substitutions of residues at equivalent positions to F279 in CC8-type FnBPB (31), CC8-type FnBPA (10), ClfA (32) and ClfB (33) resulted in a loss of fibrinogen binding. The involvement of residues N277 and F279, predicted to occupy the same positions in the binding trench of N2 and N3 as key residues involved in ligand binding by DLL in related proteins suggests that the DLL mechanism is likely involved in binding to CDSN and fibrinogen. Recently, we used single-molecule force spectroscopy to probe *S. aureus* cells expressing CC1-type FnBPB with CDSN-functionalized tips and found that the interaction involves two distinct binding sites within FnBPB (34). Each forms an extremely strong bond with CDSN. Intriguingly neither of the bonds was inhibited by a peptide representing the 17 C-terminal residues of the human Fg γ-chain (34). Here we demonstrate that CC1-type FnBPB A domain does indeed bind to the Fg γ-chain, like FnBPA and ClfA but it is possible that neither of the FnBPB binding sites are fully contained within the 17 C-terminal residues. To our surprise, incubating *S. aureus* expressing the A domain of CC1-type FnBPB with a high concentration of soluble fibrinogen prior to incubation in well coated with immobilized fibrinogen did not completely inhibit bacterial adherence to the well. The same was true when FnBPB A domain-expressing bacteria were incubated with soluble CDSN prior to their addition to plates coated in CDSN. Interestingly however we did measure some cross-inhibition of Fg binding with CDSN and vice versa. This suggests that both of these ligands bind to the same region of FnBPB. This is supported by mutagenesis studies showing that amino-acid substitutions in the ligand binding trench inhibit adhesion.

The sequences of FnBPB and ClfB are highly variable and differ between CCs while being highly conserved within CCs (35, 36). CC1 strains are the most frequently isolated from infected skin lesions in paediatric AD patients (37–39). The sequence of the well-studied CC8-type protein differs from the CC1-type FnBPB with the variation mainly occurring within the major ligand binding domains, N2 and N3 (3, 35). While our study does not directly address the contribution of sequence variation in FnBPB to ligand affinity this warrants further investigation.

The hybrid protein containing the A domain of FnBPB supported biofilm formation, presumably through facilitating homophilic interactions between FnBPB A domains on neighbouring cells (40), however, expression of the full-length FnBPB protein did not. This was surprising given that previous studies have shown that full length FnBPA and FnBPB (CC8-type) can promote biofilm accumulation. It is possible that the A domain is more highly expressed or better exposed in the hybrid protein than the full-length. In contrast to previous findings with FnBPA (18), we were able to demonstrate that mutations inhibiting ligand binding also inhibited biofilm formation suggesting that there may be differences in the mechanism of biofilm formation between FnBPA and CC1-type FnBPB. We found it particularly interesting that ligand binding and biofilm formation both appeared to be enhanced in the absence of the fibronectin binding repeat domain. Previously it has been shown that individual point mutations within the FnBRs of FnBPA have a significant impact on the binding affinity for fibrinogen (15). Perhaps the FnBRs also regulate the binding interactions of the A region of FnBPB. This warrants further investigation.

The enhancement of biofilm formation when fibrinogen was added to the growth medium was not unexpected as *S. aureus* aggregates in solutions of fibrin(ogen). This is due to the ability of Fg-binding MSCRAMMs to interact with the distal ends of the fibrinogen dimer, allowing fibrinogen to act as a linkage between neighbouring bacteria forming aggregates (41, 42). This is particularly evident in bacteria overexpressing ClfA which fail to produce biofilms in the absence of ligand, but in the presence of high concentrations of Fg biofilms are formed (Fig. 4A). Less expected was the trend towards enhanced biofilm formation when CDSN was included in the growth medium. The binding site for FnBPB in CDSN lies in the glycine serine rich region at the N-terminus of the protein which has previously been shown to be responsible for oligomerisation of CDSN (43, 44), suggesting that the binding site for FnBPB in CDSN may be distinct to that required for homo-oligomerisation. Interestingly, this region of CDSN is proteoytically removed during the terminal differentiation of the skin’s epidermidis. While the biological significance is not yet clear, it will be interesting to understand if free CDSN N-terminal region might enhance biofilm formation at the skin’s surface. Either way our work shows that the A domain of FnBPB is capable of mediating biofilm formation in vitro. Further work will focus on establishing if CC1-type FnBPB promotes biofilm formation in CC1 strains of *S. aureus*. We speculate that in the stratum corneum of AD skin, FnBPB may not only promote adherence to corneocytes via its interaction with CDSN, but also promote biofilm formation to enhance the stability of the bacteria in the stratum corneum.

## Methods

### Bacterial Strains and Plasmids

*E. coli* cloning and expression strains (Table S1) were grown in LB broth supplemented with ampicillin (100 μg/ml) as appropriate at 37°C with shaking (200 rpm). To construct the pALC2073:: *fnbB*(A) plasmid inverse PCR was performed on the pALC2073::*clfA* plasmid. Template DNA was removed by DpnI digestion. The A domain of *fnbB* was amplified from CC1 strain AD08 (3) using colony PCR with a 25 base-pair complementary overhang to the *clfA* backbone. Sequence Ligation Independent Cloning was utilised to anneal the insert and fragment. Further mutagenesis with KOD Hot Start polymerase was utilised with overlapping site directed mutagenesis primers (Table S2) to obtain the single and double amino acid substitutions at N277A and F279A. All clones were confirmed by DNA sequencing of PCR products. Subsequent to obtaining the correct clones, the plasmid DNA (Table S3) was shuffled through *E. coli* IM01B before electroporating into *S. aureus* SH1000 *clfAclfBfnbAfnbB*. Colony PCR was performed to confirm the presence of the plasmid. *S. aureus* SH1000 *clfAclfBfnbAfnbB* strains transformed with pALC2073 plasmids were subsequently grown in tryptic soy broth (TSB, Difco) supplemented with chloramphenicol (10 μg/ml) at 37°C with shaking (200 rpm).

The pQE30::*fnbB*N2N3 plasmid was used as template DNA with primers FnBPB_F1_N312A_F314A and FnBPB_R1_N312A_F314A (Table S2) to make the pQE30::*fnbB*N2N3_N277A+F279A plasmid. Subsequent to obtaining the correct clones, the plasmid DNA (Table S3) was shuffled into *E. coli* Topp3 for protein expression.

### Recombinant Protein Purification

Recombinant FnBPB was expressed and purified via nickel affinity chromatography as previously described (3, 6). Recombinant CDSNΔCT was expressed and purified by glutathione affinity chromatography as previously described (3). Protein concentration was determined by a Qubit Protein Assay Kit (ThermoFisher Scientific).

### Far-western blotting

Fibrinogen purchased from Enzymes Research Ltd, UK was separated on 10% Bis-Tris NuPAGE Gels at 180V for 75 minutes (ThermoFisher Scientific) with colourmetric protein markers (BioRad Kaleidoscope™). The gel was split and a portion was stained with SimplyBlueTM SafeStain (Invitrogen) to determine the positions of the α, β and γ chains. The other portion was transferred in 20% methanol to PVDF (1 h, 100V). The membrane was subsequently blocked (5% (w/v) BSA in PBS-T) overnight at 4°C, probed with 500 nM rFnBPB or rFnBPB_N277A/F279A in 5% (w/v) BSA PBS-T for 4 hours 20°C. The membrane was washed three times in PBS, before application of Anti-His_6_-Peroxidase (Roche, 1:1000) in 5% (v/w) BSA PBS-T. The membrane was washed 3 times in PBS, before 1 minute incubation in 1x LumiGLO®/ 1x Peroxidase (CellSignal). Blots were visualised in an Amersham A680 imager.

### Adhesion Assays

Adhesion assays were carried out as previously reported by (3), with inducer (Atc, 300 ng/ml) being added to cultures when an OD_600_ = 0.18 was reached. For ligand competition assays after adjusting the OD_600_ of the cells to 1, bacteria were incubated in the presence of no ligands, Fg, CDSN or GST for 30 min in a water bath at 37°C with regular inversion before addition to the plate. The plate was incubated statically at 37°C for 1 hour 30 minutes before washing away non-adherent bacteria and fixing adherent cells with 20% (v/v) formaldehyde.

### Biofilm Assays

4 ml overnight cultures from individual colonies were adjusted to an OD_600_ = 0.05 in fresh TSB supplemented as appropriate with chloramphenicol (10 μg/ml), anhydrotetracycline (150 ng/ml), glucose (1% w/v), and fibrinogen or CDSN at defined concentrations. Biofilms were grown statically in 200 μl volumes in 96-well plates for 24 hours. The plantonic growth and culture media was removed and the biofilm washed twice in water. Crystal violet (1% w/v) was used to stain the biofilm for minimum 1 minute before being removed and 3x washes in water to remove unbound crystal violet. The remaining crystal violet was solubilised in 70% ethanol (100 μl) and the absorbance read at 570 nm. The biofilm intensity was expressed as the absorbance relative to the biofilm produced by SH1000 *clfAclfBfnbAfnbB* pALC2073 in TSB-Cm without inducer or glucose unless otherwise stated.

### Western Immunoblotting

SH1000 *clfAclfBfnbAfnbB* strains carrying variants of the pALC2073 vector were grown overnight in TSB-Cm. The cells were washed twice in TSB before being diluted 1:100 in pre-warmed TSB-Cm and grown at 37°C 200 rpm. Cells were induced at OD_600_=0.18 with 300 ng/ml ATc. Cells were grown until OD_600_=0.35 when they were harvested and washed in equal volume of PBS. The OD_600_ was adjusted to 10. 1 ml fraction was pelleted and resuspended in EDTA-free protease inhibitor cocktail (Roche), lysostaphin (12.5 μg), 50 mM Tris-HCl (pH 7.5), 20 mM magnesium chloride and incubated for 8 minutes at 37°C. The cell debris was pelleted. The supernatent was boiled with Laemmli sample buffer (Sigma) before loading on to 10% Bis-Tris SDS-PAGE gels. Proteins were transferred to PVDF and blocked with 10% Skimmed Milk Powder in Tris-buffered saline (TBS). The membrane was subsequently probed with rabbit serum specific for the FnBPB A-domain residues 163–480 (9), 1:1000 in 10% Skimmed Milk in TBS). The membrane was washed 3 times in TBS before being probed with protein A-peroxidase (Merck, 1:50000 in 10% Skimmed Milk in TBS). The membrane was incubated with 1x LumiGLO® / 1x Peroxidase (CellSignal) for 1 minute before imaging in a GE Amersham imager AI680.

### Statistical Analysis

Statistical analysis was carried out in RStudio 3.6.3 or higher, with multcomp package 1.4-22. Significance was represented by P<0.05 *, P<0.01 **, P<0.001 ***. Typically a 1-or 2-way ANOVA was performed in combination with a post-hoc test (Dunnetts Multiple Comparison Test, where the groups are to be compared to a single reference point, or Tukey’s Honest Significant test where all points are to be compared against each other.

## Author contributions

The authors contributed to the study as follows; Conceptualization, methodology, formal analysis, writing original draft and review and editing; SRH and JAG. Investigation; SRH, Funding acquisition, project administration and resources; JAG.

## Acknowledgements

We are grateful to Giampiero Pietrocola and Pietro Speziale for providing antibodies against FnBPB. We also thank past and present members of the research group for helpful discussion.

