## Supplemental files for "The A domain of clonal complex 1-type fibronectin binding protein B promotes adherence and biofilm formation in *Staphylococcus aureus*"

**Table S1. Bacterial Strains**

| Strain | Description | Reference |
| --- | --- | --- |
| *E. coli* XL1-Blue | Host for cloning, *recA1* *endA1 gyrA96 thi-1 hsdR17 supE44 relA1 lac* | Stratagene |
| *E. coli* IM01B | Plasmid propergation. SA08BΩP_N25-_ *hsdS* (CC1-1) | (1) |
| *S. aureus* SH1000 | Derivative of strain 8325-4 with a repaired rsbU gene | (2) |
| *S. aureus* SH1000 *clfAclfBfnbAfnbB* | Adhesion deficient mutant. ClfA, ClfB, FnBPA and FnBPB are disrupted | (3) |
| *E. coli* Topp3 | Strain deficient in proteases for expression of recombinant proteins. Rifr , [F’proAB+ | Stratagene |

**Table S2. Primers**

| ClfA_BB_rev | CTGTAGTATTGTTTTGTTCCGATGCTGCATCTGCTTCTTTACTGCTGAG |
| --- | --- |
| ClfA_BB_for | TAATAACGCTCAAGGCGACGGTAAAGATTCAGATTCTGACCCAGGTTCA |
| FnBPB_N1_F | GCATCGGAACAAAACAATACTACAGTAG |
| FnBPB_N3_R | TTTACCGTCGCCTTGAGCGTTATTAGAGTA |
| FnBPB_F1_N312A_F314A | GGCTGAATTAAACTTA**GCT**TTG**GCT**ATTGATCCAAC |
| FnBPB_R1_N312A_F314A | CGTAACTGTAGTTGGATCAAT**AGC**CAA**AGC**TAAGTT |
| FnBPB_F1_N312A | GGCTGAATTAAACTTA**GCT**TTG**TTT**ATTGATCCAAC |
| FnBPB_R1_N312A | CGTAACTGTAGTTGGATCAAT**AAA**CAA**AGC**TAAGTT |
| FnBPB_F1_F314A | GGCTGAATTAAACTTA**AAT**TTG**GCT**ATTGATCCAAC |
| FnBPB_R1_F314A | CGTAACTGTAGTTGGATCAAT**AGC**CAA**ATT**TAAGTT |

Bold = Nucleotide Changes; Underline = complimentary overhangs.

**Table S3. Plasmids**

| Plasmid | Description | Reference |
| --- | --- | --- |
| pALC2073 | Tetracycline inducible expression plasmid. Cm^r^ resistance. | (4) |
| pALC2073::*fnbB* | pALC2073 carrying the *fnbB* gene from CC1 *S. aureus*. Cm^r^ | (5) |
| pALC2073::*clfA* | pALC2073 carrying the *clfA* gene from CC8 *S. aureus*. Cm^r^ | (6) |
| pALC2073::*fnbB*(A) | pALC2073 carrying the *clfA* gene from CC8 *S. aureus*, where the *clfA* A-domain is replaced with the sequence of the CC1 type *fnbB* A domain. Cm^r^ | This study |
| pALC2073::*fnbB*(A)_N277A | pALC2073 carrying the *clfA* gene from CC8 *S. aureus*, where the *clfA* A-domain is replaced with the sequence of the CC1 type *fnbB* A domain with mutations resulting in the N277A substitution. Cm^r^ | This study |
| pALC2073::*fnbB*(A)_F279A | pALC2073 carrying the *clfA* gene from CC8 *S. aureus*, where the *clfA* A-domain is replaced with the sequence of the CC1 type *fnbB* A domain with mutations resulting in the F279A substitution. Cm^r^ | This study |
| pALC2073::*fnbB*(A)_N277A+F279A | pALC2073 carrying the *clfA* gene from CC8 *S. aureus*, where the *clfA* A-domain is replaced with the sequence of the CC1 type *fnbB* A domain with mutations resulting in both N277A and F279A substitutions. Cm^r^ | This study |
| pGEX-4T2 | *E. coli* vector for the expression of glutathione S-transferase tagged recombinant proteins. Amp^r^ | GE Lifesciences |
| pGEX-4T2::CDSNΔCT | pGEX-4T2::CDSN derivative where DNA encoding residues G376-G76 have been deleted. Amp^r^ | (5) |
| pQE30::*fnbB*N2N3 | pQE30 carrying DNA encoding CC1-type FnBPB N2N3 subdomains. Amp^r^ | (7) |
| pQE30::*fnbB*N2N3_N277A+F279A | pQE30 carrying DNA encoding CC1-type FnBPB N2N3 subdomains, with mutations resulting in both N277A and F279A substitutions. Amp^r^ | This study |

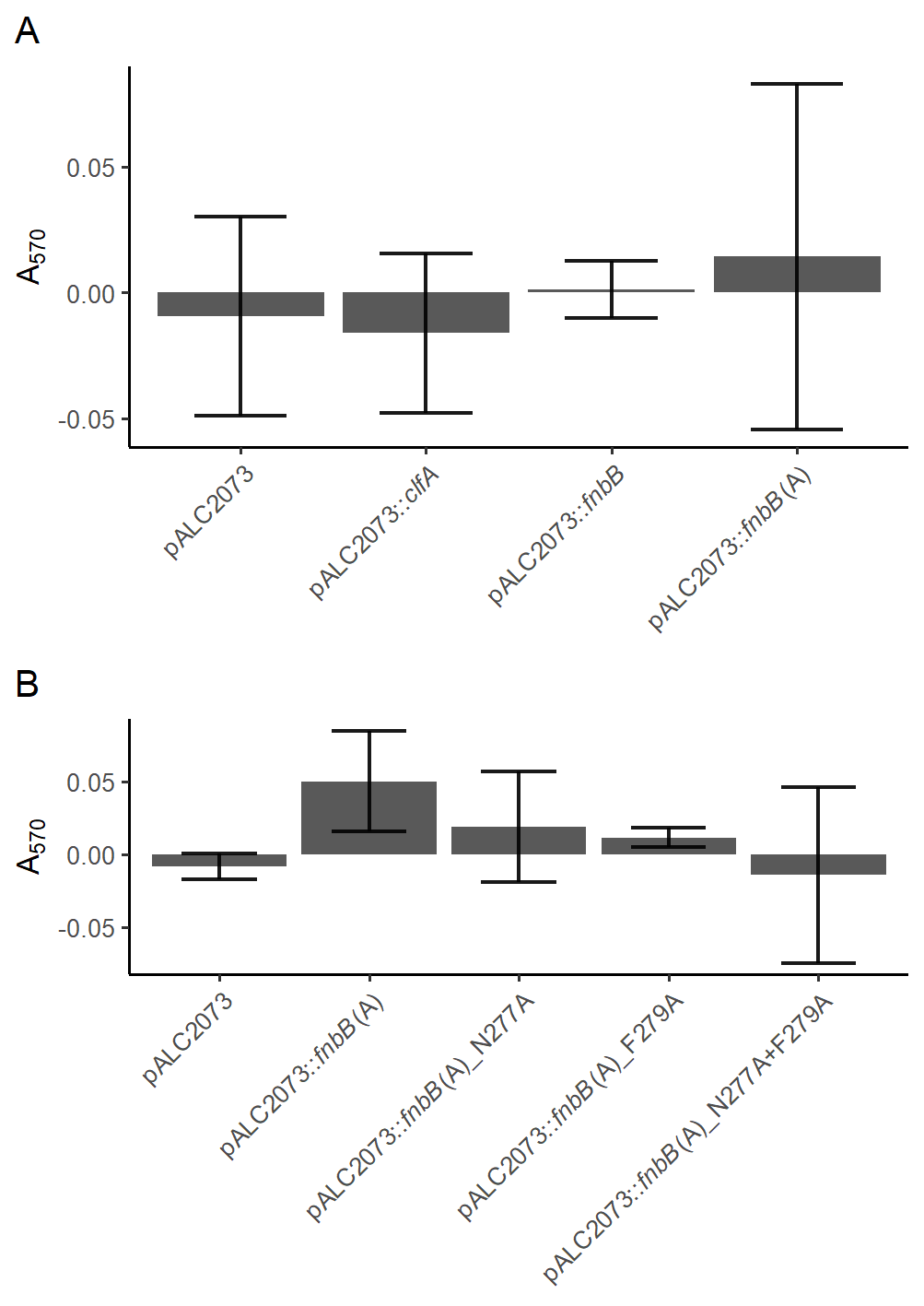

**Fig. S1:** Adhesion of SH1000 *clfAclfBfnbAfnbB* to GST (300 nM). A) Data corresponds to the strains presented in figure 1 (A) and figure 2 (B). Data is presented as the mean of a minimum of 3 biological replicates, error bars represent 1 Standard deviation from the mean. No statistical significance was determined by 1-way ANOVA with post-hoc Tukey’s honest significance test.

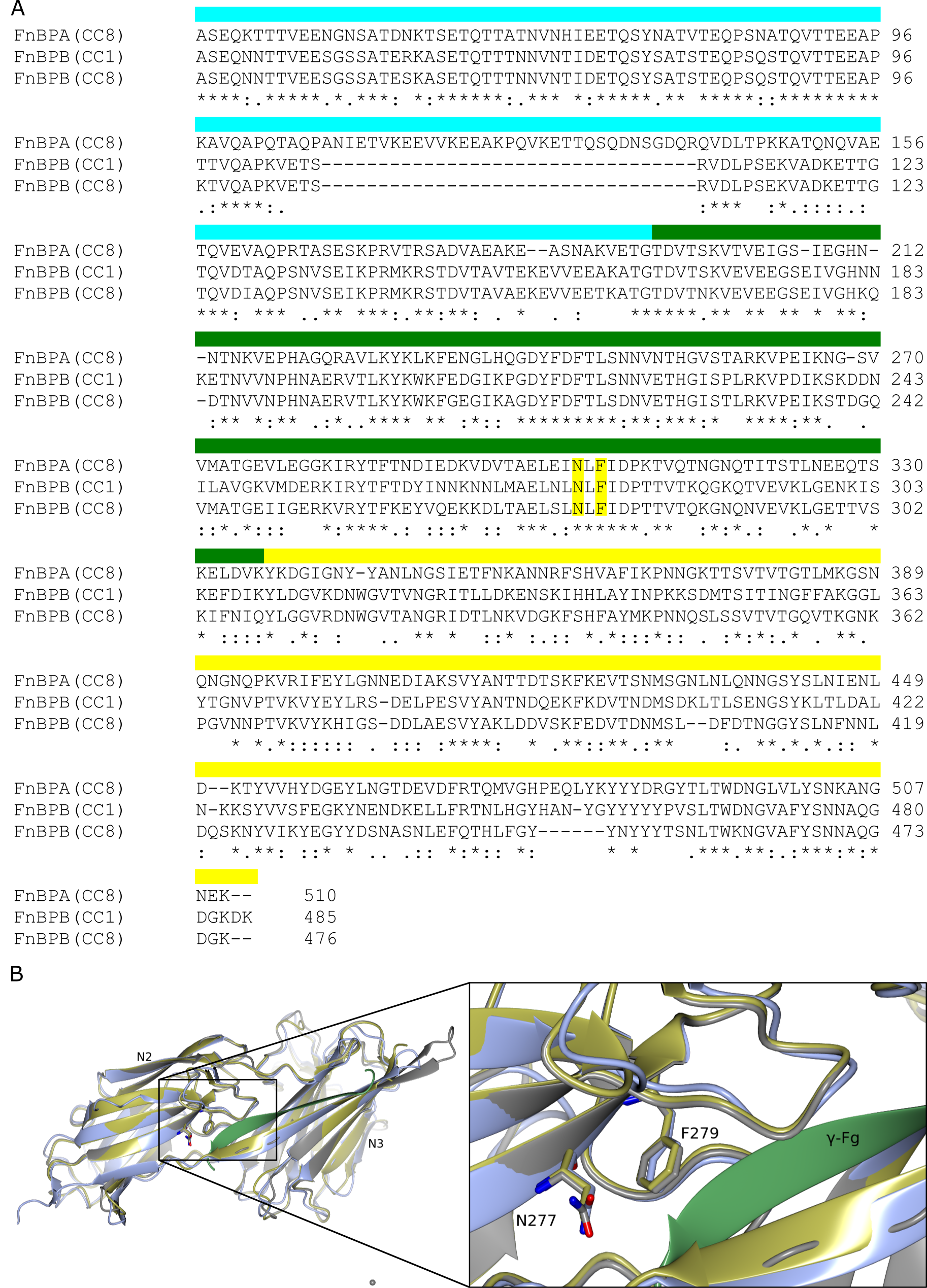

**Fig. S2.** A) Sequence alignment of the A-region comprising of the N1N2N3 domains of FnBPB from clonal complex 1 (AD08), clonal complex 8 (USA300) and FnBPA from clonal complex 8 (NCTC8325). The N1 domain is indicated in cyan, N2 in green and the N3 in yellow. Yellow highlighted residues indicate the position of the N277 and F279 residues in CC1 type FnBPB. B) Structural Alignment of the N2N3 domains of FnBPA (Gold) in the presence of the γ-Fibrinogen peptide (Green) (pdb: 4B60) aligned with the AlphaFold-2 prediction of CC8-type FnBPB (Blue) and the Phyre2 prediction of CC1-type FnBPB (Grey). Overview left and detailed view of the ligand binding trench indicating the positions of the N277 and F279 residues in relation to the predicted ligand binding site.

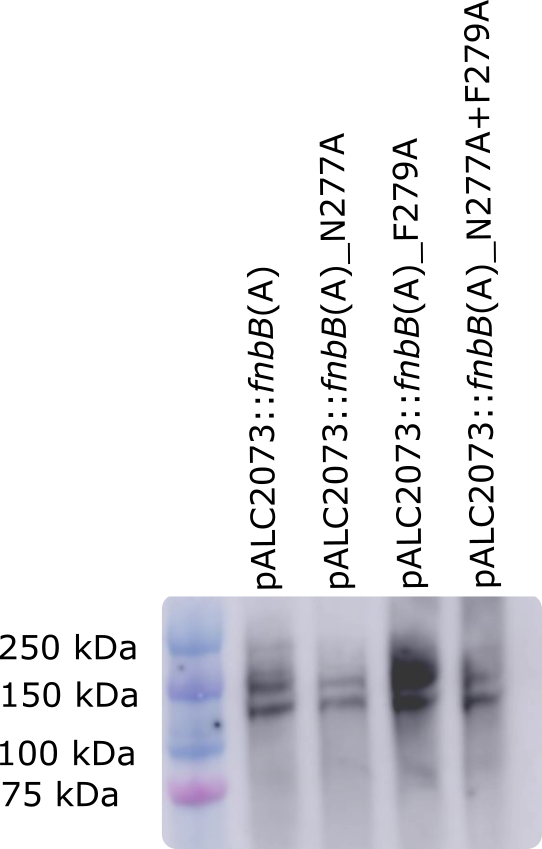

**Fig. S3:** Expression of the FnBPB A domain in SH1000 *clfAclfBfnbAfnbB* transformed with pALC2073::*fnbB*(A) variant plasmids induced with ATc (300 ng/ml) when cultures reached an OD_600_ = 0.18 and harvested at OD_600_ = 0.35. Washed cells were resuspended to an OD_600_ = 10, and the cell wall anchored proteins extracted and separated on 10% Nu-PAGE Bis-tris gels in MOPS running buffer. Proteins were transferred to PVDF membrane and subsequently blocked with 10% skimmed milk powder. Primary antibody = anti-FnBPB (1:1000). Bound antibody was detected with protein A peroxidase (1:50000).

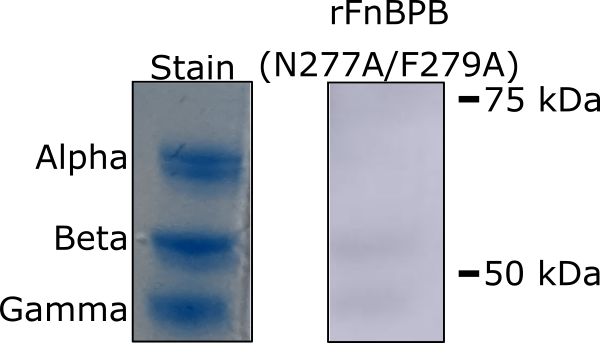

**Fig. S4:** Recombinant FnBPB N2N3 domains with N277A and F279A substitutions does not bind to the gamma chain of fibrinogen. Fibrinogen was separated on 10% Bis-tris Gels in MOPS running buffer under reducing conditions. The stained portion of the gel was stained to distinguish the positions of the alpha, beta and gamma Fibrinogen chains in decreasing molecular mass. The rFnBPB portion of the gel was transferred to PVDF membrane, blocked with 5% (w/v) BSA (PBS-T) overnight (4°C) and probed with recombinant rFnBPB for 4 hours at room temperature. Bound rFnBPB was detected with HRP-conjugated Anti-His antibody (Roche, 1:1000 in 5% (w/v) BSA PBS-T), and incubated with LumiGlo reagent before chemiluminescent imaging.

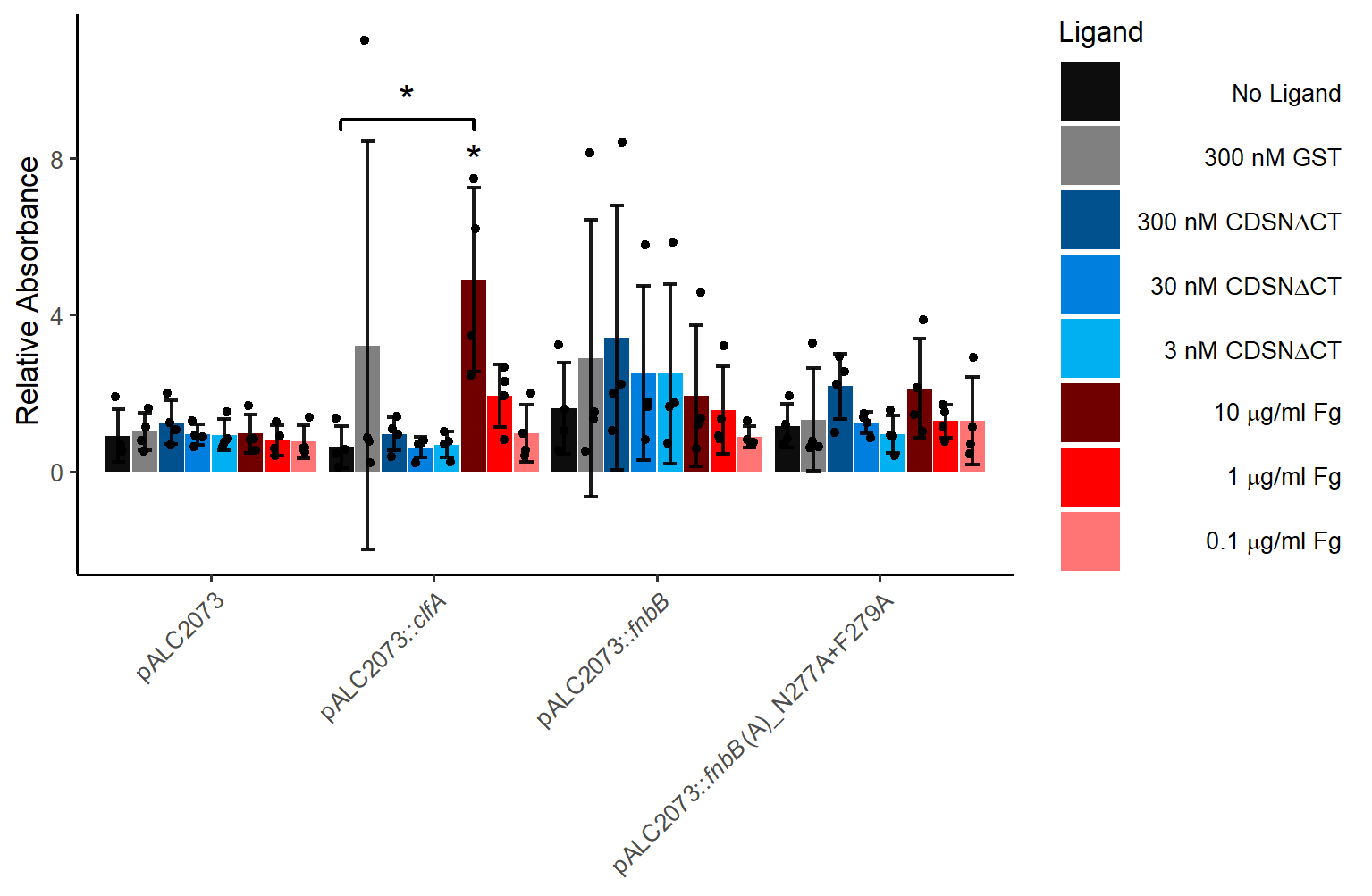

**Fig. S5.** The impact of FnBPB ligands on biofilm formation. 24 hour biofilms of SH1000 *clfAclfBfnbAfnbB* transformed with pALC2073 and derivatives were grown in TSBG supplemented with anhydrotetracycline (150 ng/ml) with ligands at the specified concentrations. Biofilms were washed and subsequently quantified by staining with crystal violet (1% w/v) solubilised in 70% ethanol. To determine the relative absorbance values the A_570_ values for each biological replicate were divided by those obtained in the absence of a ligand. Data is presented as the mean of 4 biological replicates. Error bars represent 1 standard deviation from the mean. Significance was tested via a 1-way ANOVA test with a Dunnett’s Multiple Comparison test comparing each strain to the biofilm formed in the absence of any ligand. *p<0.05.
